# RNA-seq analysis of transcriptomes for assessing stress tolerance of *S. cerevisiae* strain, NCIM3186

**DOI:** 10.1101/609370

**Authors:** Burragoni Sravanthi Goud, Kandasamy Ulaganathan

## Abstract

We have previously sequenced the draft genome of high ethanol producing *S. cerevisiae* strain, NCIM3186. Towards assessing the stress tolerance by this strain transcriptomes from control and in response to glucose, ethanol and furfural stress were sequenced. Comparative RNA-seq analysis of these transcriptomes identified 573 differentially expressed genes of which thiamine biosynthesis genes under furfural stress, TDH1, heat shock proteins and hexose transporter gene under ethanol stress were observed to be highly differentially expressed. Apart from thiamine biosynthesis genes and TDH1, 2 other proteins of unknown function were highly differentially expressed under glucose stress. Most importantly, TAR1 gene was highly down-regulated under all the stress conditions compared to control. Among 93 fermentome genes, 7 (TPS1, TPS2, SIN3, PTK2, SSQ1, ZAP1, DOA4) out of 9 stuck genes are found to be differentially expressed. Several stress-related genes like PHO4, SOD2, STR3, GRE2, GLR1, MEP1,3, MLH3, SNF1, MSN2, ATG1, GLC7 were differentially expressed.

## 1. Introduction

Reduction in fossil fuel consumption by using alternate sources of energy is the major challenge to be addressed in the coming decades. Bioethanol is considered to be the most viable option for addressing this challenge. Lignocellulosic biomass is the best source of bioethanol production. Ample quantities of lignocellulosic biomass (60 billion tons) are available from terrestrial plants [1] which is renewable, and can be used without disturbing the food, economy and the environment [2,3,4]. Economically viable bioethanol production from lignocellulosic biomass is yet to be realized due to the existence of several barriers [5]. Efforts are underway to break the barriers utilizing the unprecedented tools made available by the genomic revolution sweeping biology recently. Precision genome engineering is the latest among the tools contributed by the field of genomics [6].

An ideal organism for lignocellulosic bioethanol production should have the following characters: utilize lignocellulose, ferment hexose and pentose sugars, high ethanol yield, tolerate high ethanol concentration, higher temperature and extreme pH, amenable for genetic manipulation, availability of recombinant DNA methods for modification and introduction of genes suitable for heterologous expression of proteins [7,5]. Two alternate strategies are usually employed for developing a host organism for industrial lignocellulosic bioethanol production. The first one is screening different species capable of lignocellulosic bioethanol production to identify a particular species based on its performance and genetically improve it. The alternate strategy employed is to select a species like *Saccharomyces cerevisiae* which is already employed in bioethanol production and carry out targeted strain optimization. *S. cerevisiae* is the most widely used organism as it meets most needs of the bioethanol production process and its inability to utilize pentose sugars has been addressed by genetic engineering [8,9]. Owing to its ability in fermentative production of high ethanol, inhibitor tolerance, and suitability for heterologous expression of genes *S. cerevisiae* is highly preferred [5,10,11].

Understanding the genomic variations that facilitate high ethanol production by *S. cerevisiae* is necessary for engineering strains for lignocellulosic bioethanol production. Many strains used in bioethanol production have been sequenced, and a number of variations have been identified [6]. In our effort to select a suitable strain for lignocellulosic bioethanol production, we have sequenced strains differing in their ability to produce bioethanol from plant biomass and reported the genome sequences of a moderate and high ethanol producing strains NCIM3107 and NCIM3186, respectively [12,13,14].

Stress tolerance mechanisms in *S.cerevisiae* are highly diversified depending upon the stress conditions posed to it. For a bioethanol producing yeast strain ethanol, inhibitors (from lignocellulose biomass), thermal, acid and nutrient stress conditions are the major challenges posed at industrial scale [15]. In this study, we aimed at understanding the gene expression pattern of high ethanol producing yeast strain, NCIM3186 under ethanol, furfural (inhibitor), glucose stress conditions. We sequenced transcriptomes of control and stress treated NCIM3186 strain and carried out comparative RNA-seq analysis and the results are reported here.

## 2. Materials and Methods

### 2.1. Strain and culture conditions

The yeast strain used in this study is *Saccharomyces cerevisiae* NCIM3186 strain, collected from the Microbial Type Culture Collection, Chandigarh, India in the form of lyophilised powder. The obtained culture was then revived according to the MTCC prescribed protocol using distilled water. YEPD medium composed of yeast extract (0.3%), peptone (1%), glucose (2%) [for broth cultures] and agar (1.5%) [for plate cultures] was used for growing the yeast cultures. For stress treated samples growth YEPD medium + 8%(v/v) ethanol for ethanol stress, YEPD medium + 1%(g/l) furfural for furfural stress, YEPD medium with 4% glucose for excess-glucose and regular YEPD medium as common control were used.

### 2.2. Sample preparation and RNA isolation

Yeast pre-culture was prepared by inoculating a pure single colony into fresh YEPD broth and incubated at 30°C for 24hr without shaking. After 24hr incubation, 1% (v/v) yeast pre-culture inoculum was collected in 4 centrifuge tubes & centrifuged at 6000 xg for 5 min. Supernatant was discarded and the pellets were re-suspended in 1ml of ddH_2_0. 1ml each of the above culture was added to 4 culture flasks with screw caps which are incubated overnight under anaerobic condition until the culture reached to an OD_600_ value of 0.85-0.95. Of these, 1st flask contained YPD with 2% glucose which is used as common control (Control) and 2nd one contained YPD with 4% glucose which is Glucose-stressed one (Glucose). These 2 samples were collected and centrifuged at 6000 xg for 5 min. Supernatant was discarded, pellet was washed with water, RNAlater added and frozen in liquid nitrogen and used for RNA isolation and sequencing. The 3rd flask containing YPD with 2% glucose was treated with 8% (v/v) ethanol for 2hrs which is ethanol-stress sample (Ethanol) and the last flask containing YPD with 2% glucose was treated with 1% (g/l) furfural for 4hrs which is furfural-stress sample (Furfural) and samples were collected and stored for RNA isolation. Pure RNA was extracted using HiPurA Yeast RNA isolation kit method. RNA quantification was performed with Qubit 2.0 (Life Technologies, Carlsbad, CA, USA). The quantity and integrity of the extracted RNA was determined using a NanoDrop ND-1000 spectrophotometer (Nanodrop Technologies, Wilmington, USA) and by electrophoresis on 1.2 % agarose gel.

### 2.3. Library preparation and RNA sequencing

Library preparation was done using Illumina TruSeq RNA library protocol developed by Illumina Technologies (San Diego, CA). 1 ug of total RNA was subjected to PolyA purification of mRNA. Purified mRNA was fragmented for 8 minutes at elevated temperature (94°C) in the presence of divalent cations and reverse transcribed with SuperScript III Reverse Transcriptase by priming with random hexamers. Second strand cDNA was synthesized in the presence of DNA polymerase I and RnaseH. The cDNA was cleaned up using HighPrep PCR (MAGBIO, Cat# AC-60050). Illumina adapters were ligated to the cDNA molecules after end repair and addition of A base. SPRI (solid-phase reversible immobilization, Beckman Coulter) cleanup was performed after ligation. The library was amplified using 8 cycles of PCR for enrichment of adapter ligated fragments. The prepared library was quantified using Qubit and validated for quality by running an aliquot (1 μl) on High Sensitivity DNA Kit (Agilent) which showed expected fragment distribution in the range of ~250–500 bp. The effective sequencing insert size was ~130–380 bp; the inserts were flanked by adapters whose combined size was ~130 bp. Transcriptome sequencing was carried out with the Illumina Hiseq 2500 system (Illumina, San Diego, CA) at Agrigenomes lab facility.

### 2.4. Bioinformatic analysis of transcriptome data

Paired-end reads generated by RNA-seq were subjected to a round of quality trimming using Cutadapt [16] to obtain clean reads. Quality assessment report of these reads were then obtained using FastQC tool. De novo assembly of trimmed reads was performed using Trinity [17] assembler. Differential expression profiling was done by EdgeR [18] (with P_value_=1e-3, foldchage C=2) and corresponding heatmaps were generated using Clustvis online tool [19], respectively. Variation and alternate splicing events finding were called using kissplice2reftranscriptome tool [20]. To find coding and non-coding genes, families, transmembrane domains, repeats transcriptome annotation was performed using Blast2go tool [21]. KEGG pathway enrichment analysis was also completed by KAAS server [22].

### 2.5. Quantitation of transcriptome expression and DEGs identification

All the 4 samples were aligned using Bowtie2 to their respective whole transcriptome with TPM (transcripts per million) and FPKM (Fragments per Kilobase of Exon per Million Fragments Mapped) intervals. The expected counts were produced by RSEM [23] perl script, align_and_estimate_abundance.pl which comes as part of the Trinity software. The expression matrices were then computed. Normalization of FPKM values was done by using Trimmed mean of M-values (TMM) normalization present in EdgeR package. Isoform-level transcript matrices obtained by RSEM were used by EdgeR (P_value_=1e-3, fold change C=2) to identify differentially expressed genes through analyze_diff_expr.pl perl script.

### 2.6. qRT-PCR

RNA isolation was done by using HiPurA Yeast RNA isolation kit protocol. To validate the identified DEGs qRT-PCR was performed using Applied Biosystems 7500 Fast qRT-PCR machine according to the manufacturer’s protocol keeping GAPDH as the reference gene. Primers for selected DEGs were designed by using Primer3-blast as shown in Table 1 [24] and annealing temperatures were optimized for each gene successfully.

**Table 1.**
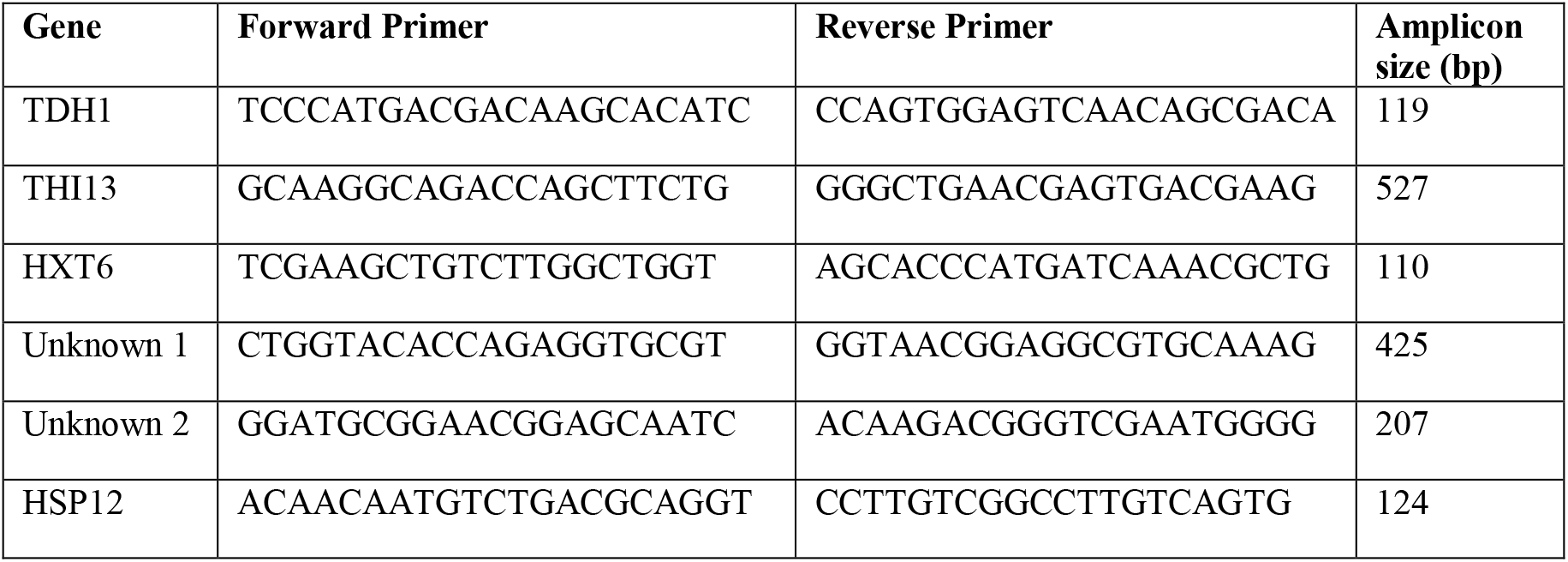
Primers used for RT-PCR validation of differentially expressed genes in NCIM3186.

## 3. Results and Discussion

### 3.1. RNA isolation, library preparation and sequencing

A fine quality intact RNA absolutely free of contamination was extracted with high yield from control and stressed (glucose, ethanol, furfural) cells of NCIM3186 strain (RIN values −8.9 to 9.5). Any traces of DNA contamination observed was removed by on-column DNase digestion. Sequencing libraries were constructed using Illumina TruSeq RNA library protocol which includes reverse transcription by SuperScript III Reverse Transcriptase followed by HighPrep PCR adapter ligation and SPRI (solid-phase reversible immobilization, Beckman Coulter) cleanup. Transcriptome sequencing was performed by using Illumina Hiseq sequencing platform (Illumina, San Diego, CA).

### 3.2. Pre-processing and de novo assembly

Transcriptome sequencing generated 22.9 - 47.2 million paired-end reads per sample with 60-100x coverage as shown in Table 4. Pre-processing of the sequenced raw reads was performed to remove any adapter contamination from the reads. Trinity based de novo assembly of these pre-processed reads generated a total of 17133 transcripts, belonging to 15103 loci with 38.6% GC content as given in Table 2. Variation analysis of the transcriptomes reported a number of short indels, single nucleotide variations, inexact tandem repeats and others which are summarized in Table 3. The paired-end reads of the sequenced transcriptomes and the transcripts of each sample were submitted to SRA and TSA, respectively under NCBI. Transcriptomes read length was 100bp. A Bioproject was created in NCBI with ID PRJNA434499 under which 4 individual Biosamples were created for control, glucose, ethanol and furfural treated transcriptomes details of which are provided in Table 4.

**Table 2.** De novo assembly statistics

| Assembly parameter | Trinity transcripts |
| --- | --- |
| Total no of transcripts | 17133 |
| Percent GC (%) | 38.60 |
| Contig N50 (bp) | 3982 |
| Contig N75 (bp) | 2322 |
| Contig L50 | 1634 |
| Contig L75 | 3511 |
| Median contig length (bp) | 544 |
| Average contig length (bp) | 1564.38 |
| Largest contig length (bp) | 38319 |
| Smallest contig length (bp) | 201 |
| Total assembled bases | 26802480 |

**Table 3.** Summary of variations identified in transcriptomes

| Type of variation | Control | Glucose | Ethanol | Furfural |
| --- | --- | --- | --- | --- |
| Single SNPs | 29509 | 26761 | 29652 | 27816 |
| Inexact tandem repeats | 31 | 33 | 34 | 28 |
| Short indels (<3nt) | 4837 | 3023 | 4304 | 3572 |
| Others | 333 | 210 | 324 | 241 |
| Alternative splicing events | 625 | 438 | 613 | 466 |

**Table 4.**
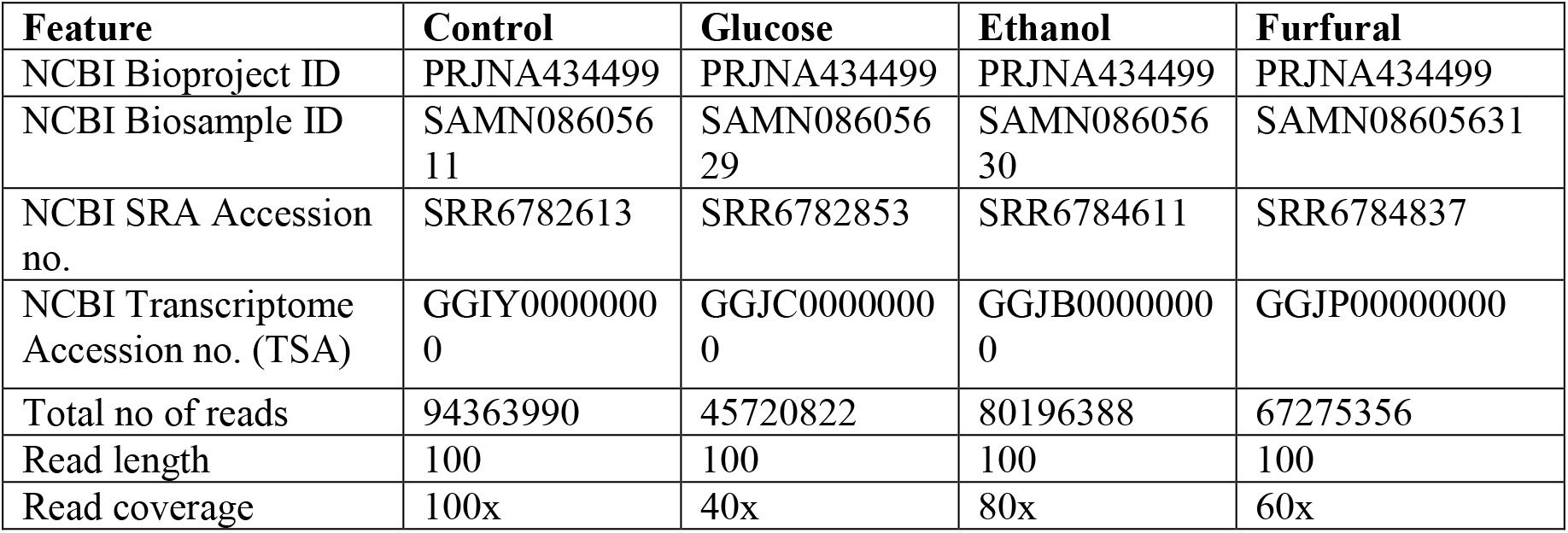
Details of transcriptomes sequenced

### 3.3. Gene Expression profiling

#### 3.3.1. Global gene expression in response to stress

Yeast cells when treated with several stress conditions showed diverse expression patterns. Under all the stressed conditions and control, a total of 15133 transcripts excluding isoforms were found. Several enzymes, transporters, transcription and translation factors, stress-related genes and most importantly fermentome genes showed significant expression levels. Fig. 1 shows that glyceraldehyde-3-phosphate genes TDH1, TDH2, TDH3, thiamine biosynthesis genes THI13, THI4, cell wall mannoprotein CCW12, stress-related gene TAR1, pyruvate kinase CDC19, snoRNA SNR37, snRNA LSR1, ribosomal proteins P2B, Rpl10, nuclear RNA TPA and non-coding RNA SCR1 were the top 15 highly expressed genes across these transcriptomes. The most striking feature is the high expression of non-coding RNA, SCR1 and other non-coding RNAs above all the coding genes.

**Fig. 1.**
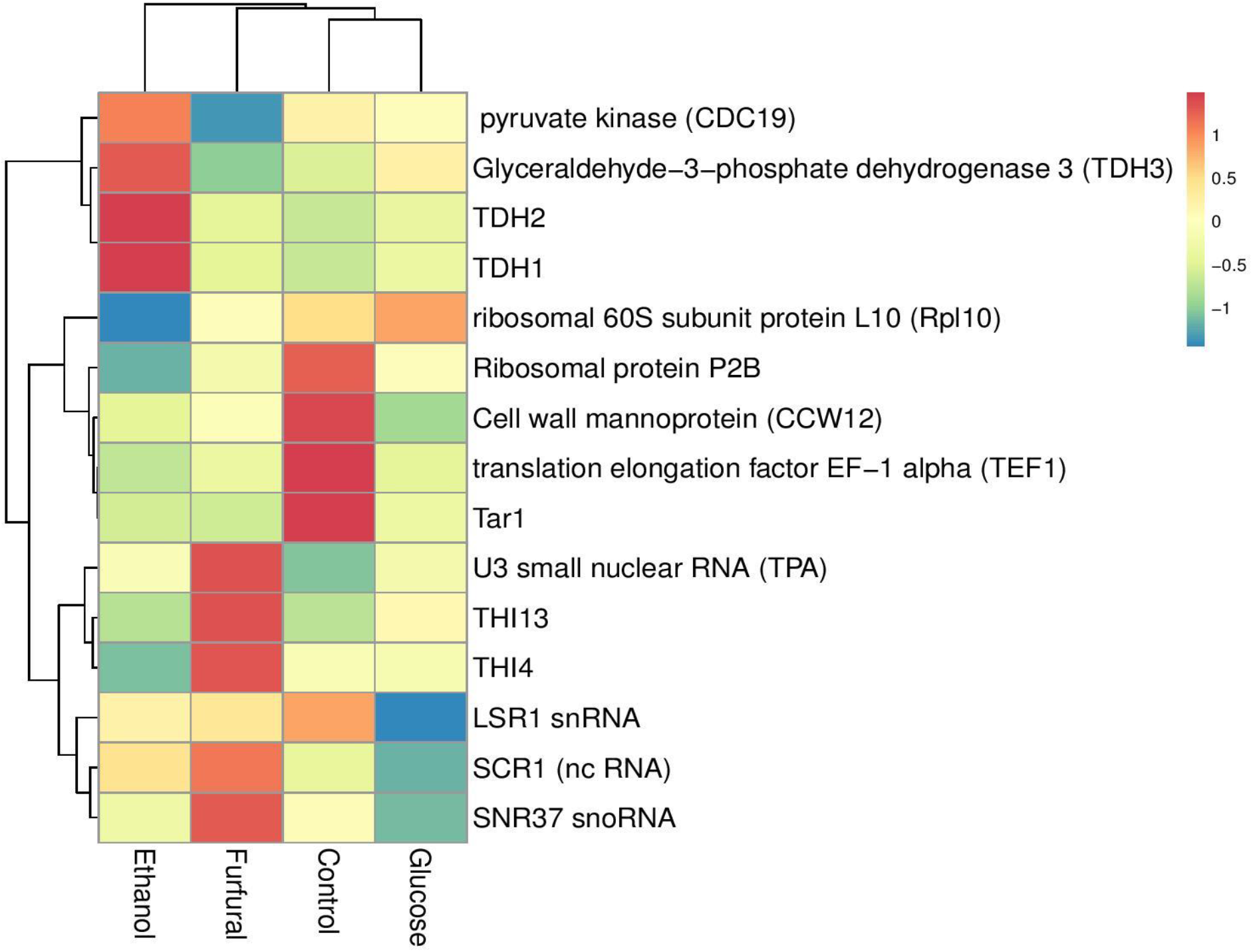
Highly expressed genes across the stress-treated transcriptomes with respect to the common control transcriptome. SCR1, a non-coding RNA topped the expression level followed by TPA, TEF1, SNR37 and TAR1.

SCR1 is an abundantly expressed small cytoplasmic RNA predominantly present in cytoplasm which mediates the translocation of membrane and secretory proteins into the endoplasmic reticulum [25]. It is the 7SL RNA subunit of SRP (Signal Recognition Particle) which is neither 3′-polyadenylated nor 5′-trimethylguanosine capped. This RNA also plays an important role in maintaining normal growth, cell division, and mitochondrial stability [26]. In our study, high expression of this particular small RNA reflects its possible regulatory role in yeast under stress conditions. Apart from this, high expression of other non-coding rRNAs, sno and snRNAs strongly supports the fact that though non-coding RNAs cannot produce functional proteins, their regulatory role and involvement in altering the expression of coding genes is highly crucial.

#### 3.3.2. Expression of Fermentome genes

“Fermentome” is a set of 93 genes in a laboratory yeast which are very much required for the timely completion of the fermentation process. Deletion or loss of function of the 9 genes (TPS1, TPS2, SIN3, PTK2, SSQ1, ZAP1, DOA4, NPT1, PLC1) named “stuck genes” out of these 93 would result in the complete cessation of the fermentation called stuck fermentation [27]. Deletion or loss of function of the remaining 84 genes named “protracted genes” would lead to the retarded fermentation called protracted fermentation. In our study, we looked at the expression of the stuck genes in NCIM3186 which showed that only 7 out of 9 stuck genes were differentially expressed across the 4 samples of which TPS1 was highly differentially expressed followed by TPS2, SIN3, PTK2, SSQ1, ZAP1, DOA4 (Fig. 2). TPS1 and TPS2 code for Trehalose 6-phosphate synthase and phosphatase respectively, both of which synthesize the storage carbohydrate trehalose and their expression is induced by the stress response [28]. Overexpression of TPS1 and TPS2 genes lead to enhanced thermotolerance in yeast during ethanol fermentation [29]. SIN3 codes for transcription co-factor subunit of Rpd3S and Rpd3L histone deacetylase complexes involved in transcriptional repression and maintenance of chromosomal integrity [30]. PTK2 is a Serine/threonine protein kinase involved in regulation of ion transport across plasma membrane [31]. SSQ1 is a mitochondrial hsp70-type molecular chaperone belonging to stress seventy subfamily Q and required for assembly of iron/sulfur clusters into proteins at a step after cluster synthesis and for maturation of Yfh1p [32]. ZAP1 is a zinc-regulated transcription factor having seven zinc finger domains which binds to zinc-responsive promoters to induce transcription of certain genes [33]. DOA4 is a ubiquitin hydrolase that de-ubiquitinates intralumenal vesicle (ILVs) cargo proteins and also required for recycling ubiquitin from proteasome-bound ubiquitinated intermediates [34].

**Fig. 2.**
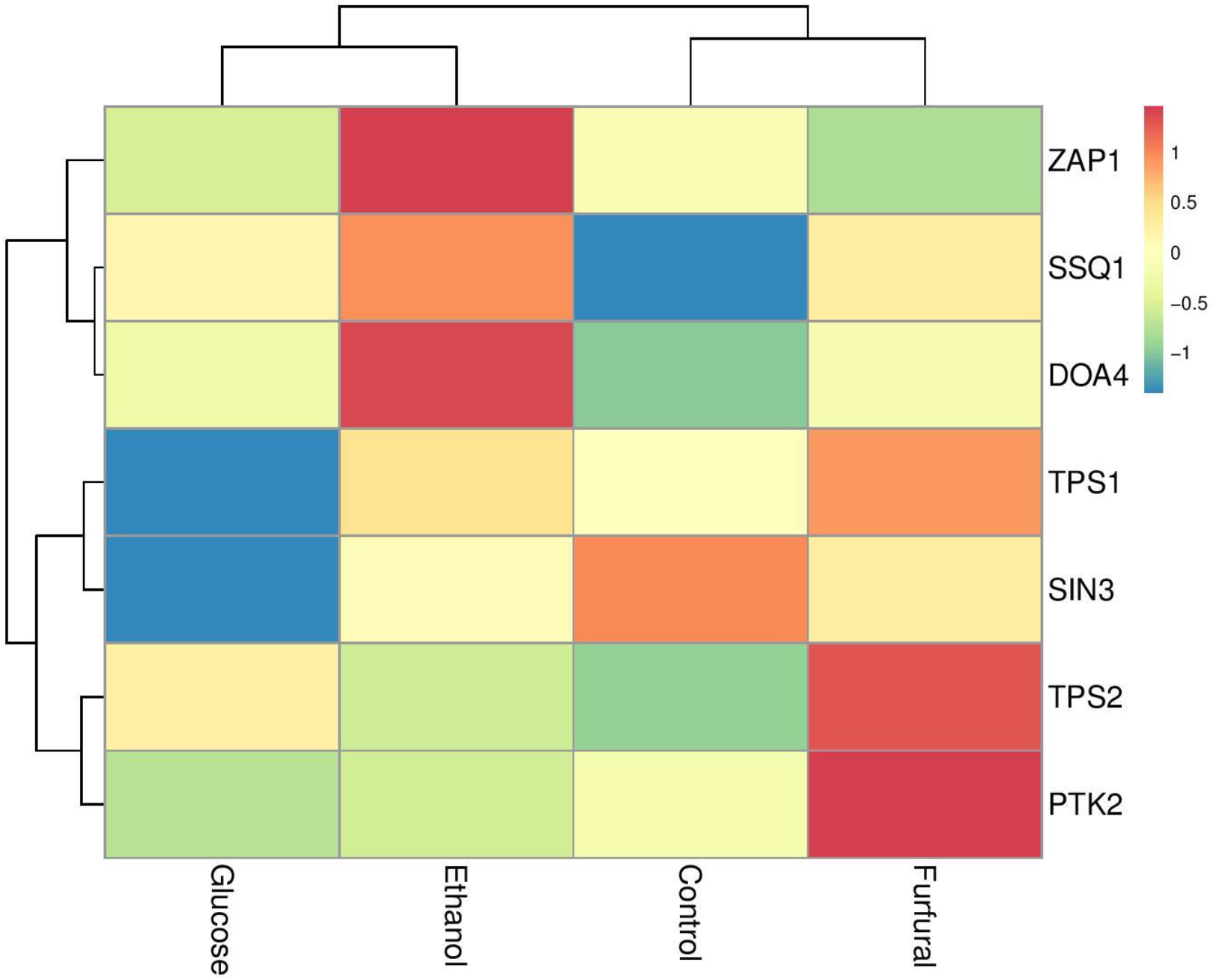
Expression of fermentome genes under stress conditions in NCIM3186. Of 93 fermentome genes, 7 out of 9 stuck genes were differentially expressed. TPS1 and TPS2 genes were highly differentially expressed.

### 3.4. Differential Gene Expression

A total of 573 DEGs were identified by differential expression profiling across 3 different conditions along with the common control. When compared to the control separately, 204, 305 and 210 genes were differentially expressed in ethanol, furfural and glucose treated cells, respectively. Several important transporters, transcription and translation factors and a large number of different enzymes were found to be differentially expressed across the transcriptomes as depicted in Fig. 3,4,5 respectively.

**Fig. 3.**
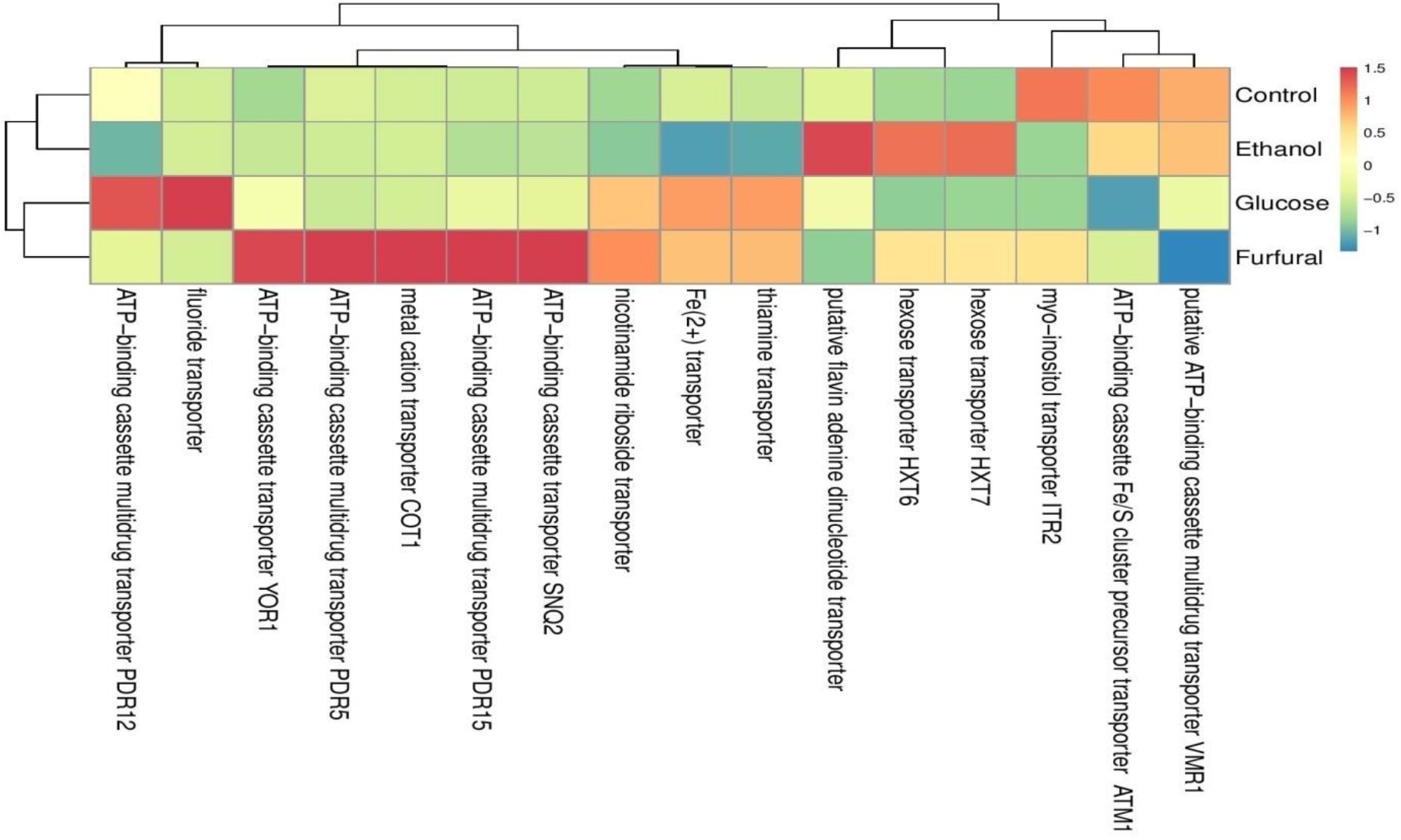
Differential expression of transporter genes in the stress treated transcriptomes of NCIM3186. HXT7, HXT6, SNQ2, PDR5 and a thiamine transporter genes have shown significant differential expression.

**Fig. 4.**
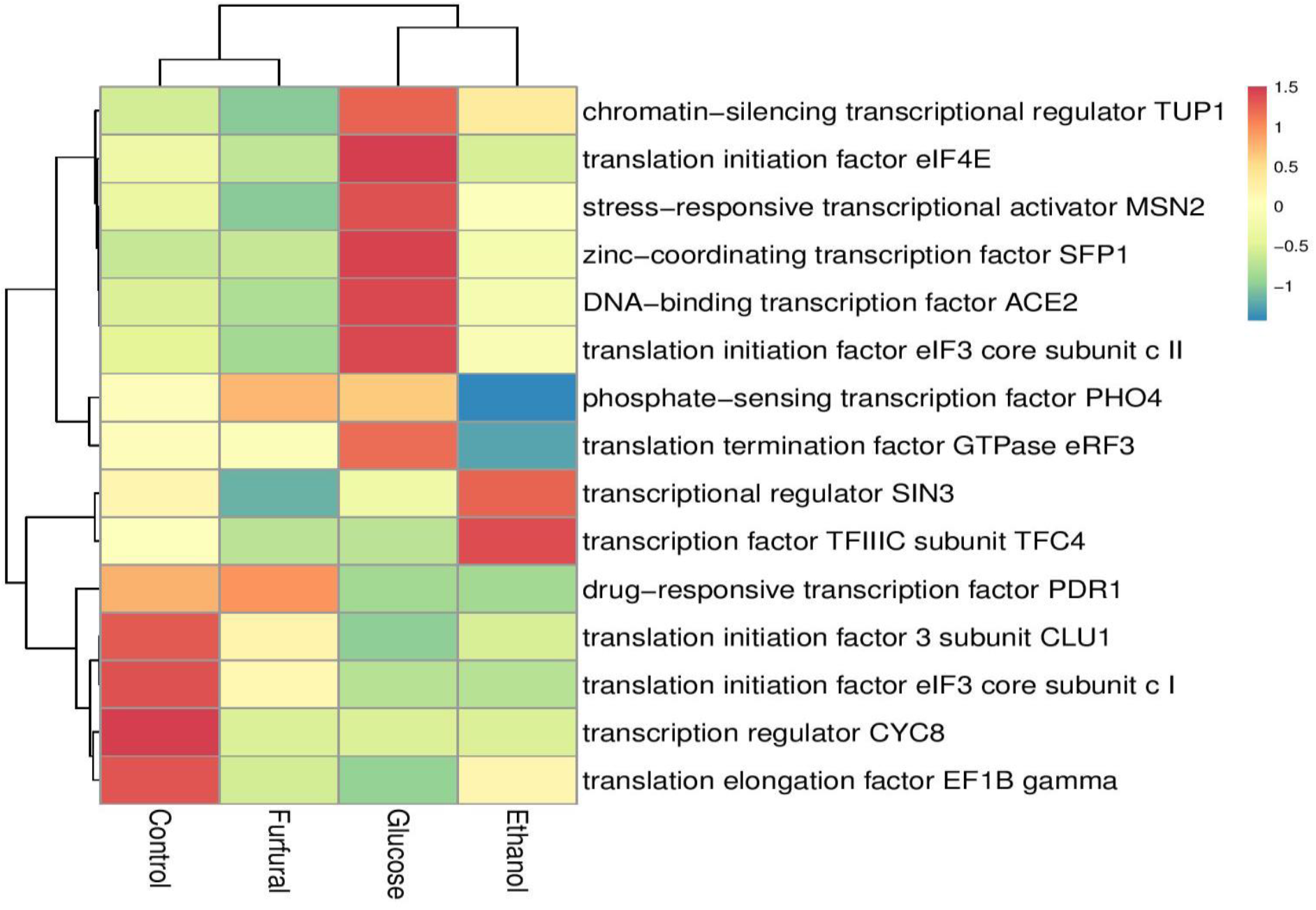
Differential expression of transcription and translation factor genes in the stress treated transcriptomes of NCIM3186. Translation termination factor GTPase eRF3, translation initiation factor eIF3 and a phosphate sensing transcription factor PHO4 genes showed significant differential expression.

**Fig. 5.**
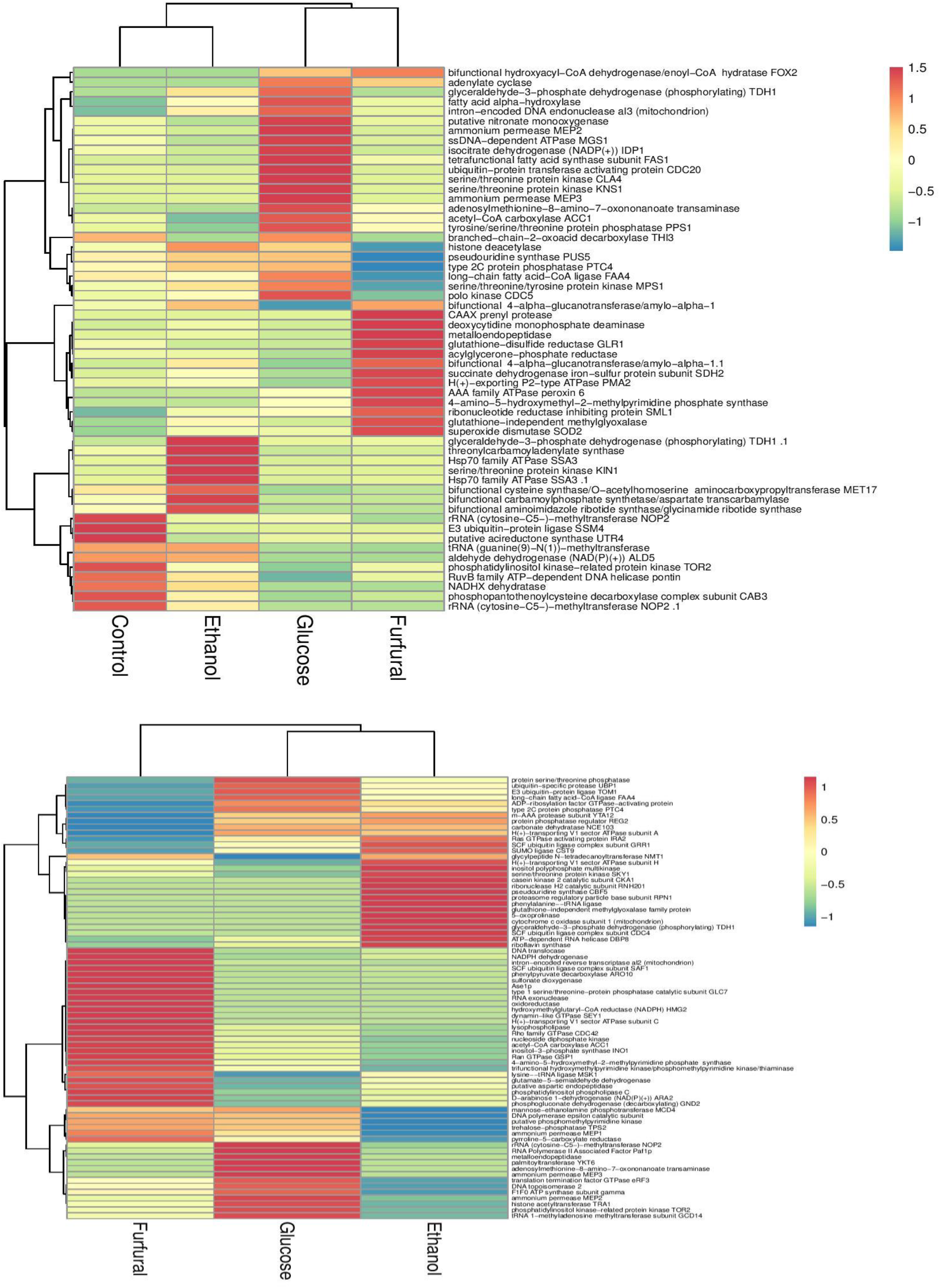
Differential expression of enzyme coding genes of NCIM3186 under stress conditions. TDH1, STR3, GRE2, peroxin 6, MEP1, ACC1, GLR1, eRF3 genes were highly differentially expressed.

#### 3.4.1. Highly Differentially expressed genes

Among 573 DEGs, top 20 highly differentially expressed genes were glyceralehyde-3-phosphate dehydrogenase TDH1, thiamine synthase THI13, hexose transporter HXT6, pleiotropic drug resistance transporter PDR5, heat shock proteins HSP26 and HSP12, STR3, INO1, TAR1, SSA3, MNT3, PEX6, RGI1, IRC8, VMA13, FAA4, YRO2, OLI1, intron-encoded reverse transcriptase al2 and 2 unknown proteins as shown in Fig. 6. This implies that in NCIM3186, under stress conditions, glycolysis regulatory enzyme TDH1, thiamine metabolism gene, certain crucial sugar and multidrug transporters, heat shock proteins, anti-oxidant enzymes, few stress related genes and some of the stress-induced proteins of unknown function altogether serve as a repertoire of yeast stress tolerance by means of their altered expression.

**Fig. 6.**
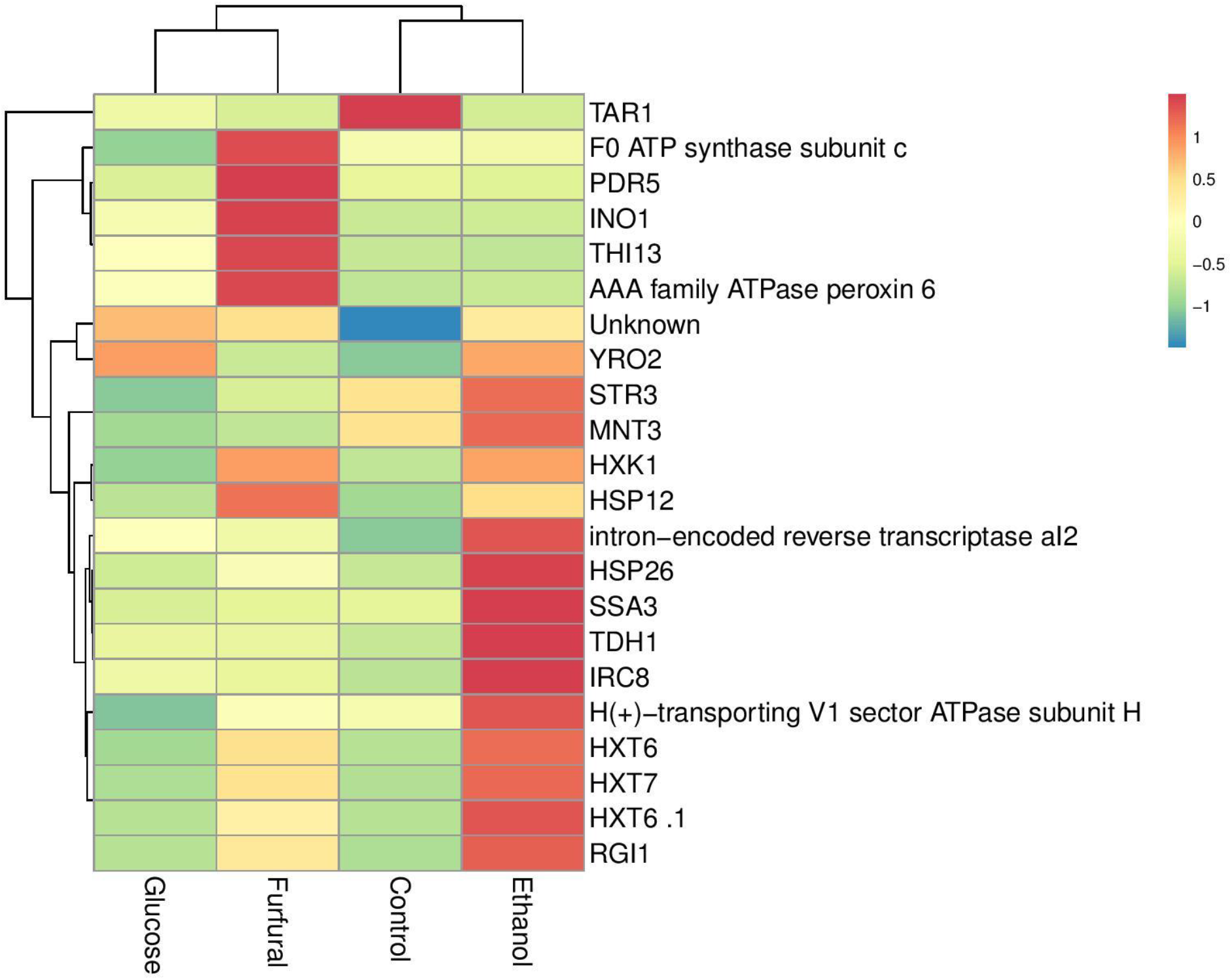
Genes showing high level of differential expression in NCIM3186 under stress conditions. TAR1, THI13, TDH1, HSP26 were highly differentially expressed genes among 573 DEGs.

#### 3.4.2. Thiamine biosynthesis genes are highly up regulated under furfural stress

Thiamine is a water soluble B-vitamin which is very essential for fermentation of sugar, defense against oxidative and osmotic stress in *S. cerevisiae*. Though only few reports suggest a relationship between thiamine and yeast cellular stress responses, there exists an important regulatory role for thiamine under stress [35,36,37]. Under stress conditions, yeast accumulates free thiamine which implies the protective role of thiamine in *S. cerevisiae* [38]. Activation of thiamine biosynthesis is a way of compensating the stress response disruption. In our study, THI13, a member of the *THI5* family (*THI5/11/12/13*) showed increased expression levels under furfural stress followed by glucose and ethanol stress which confirms the role of thiamine in yeast stress response. Not only THI13 but also other thiamine biosynthetic pathway genes like THI2, THI3, THI20, THI22, THI74 repressible mitochondrial transporter and a thiamine transporter also showed significant differential expression which is clearly shown in Fig. 7,8.

**Fig. 7.**
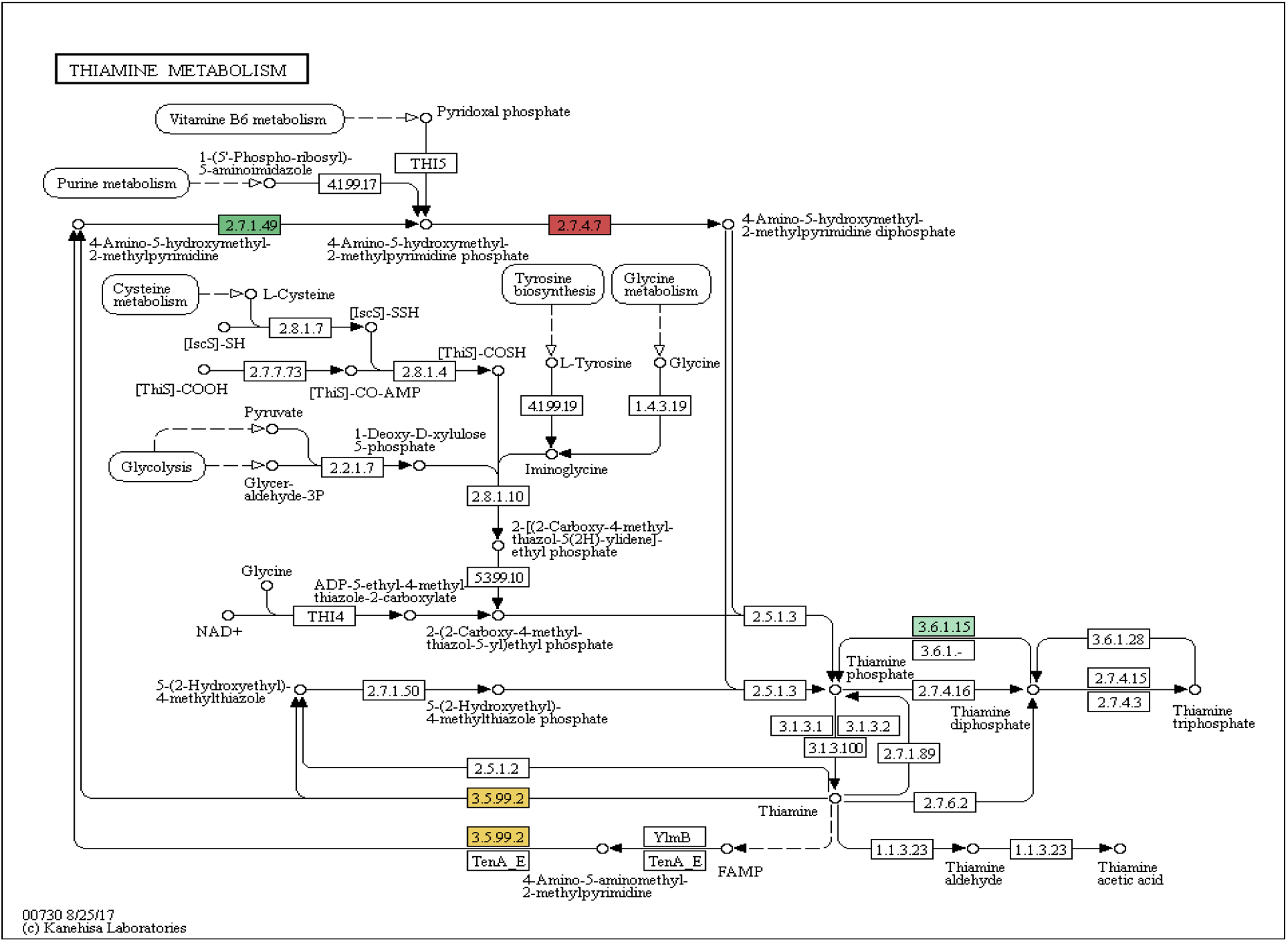
Thiamine biosynthesis pathway (KEGG) showing differentially expressed thiamine biosynthesis genes in stress treated transcriptomes of NCIM3186. Thiamine biosynthesis genes coding for enzymes with EC number 2.7.1.49, 2.7.4.7, 3.6.1.15 and 3.5.99.2 were coloured differently showing that these genes are differentially expressed under furfural stress.

**Fig. 8.**
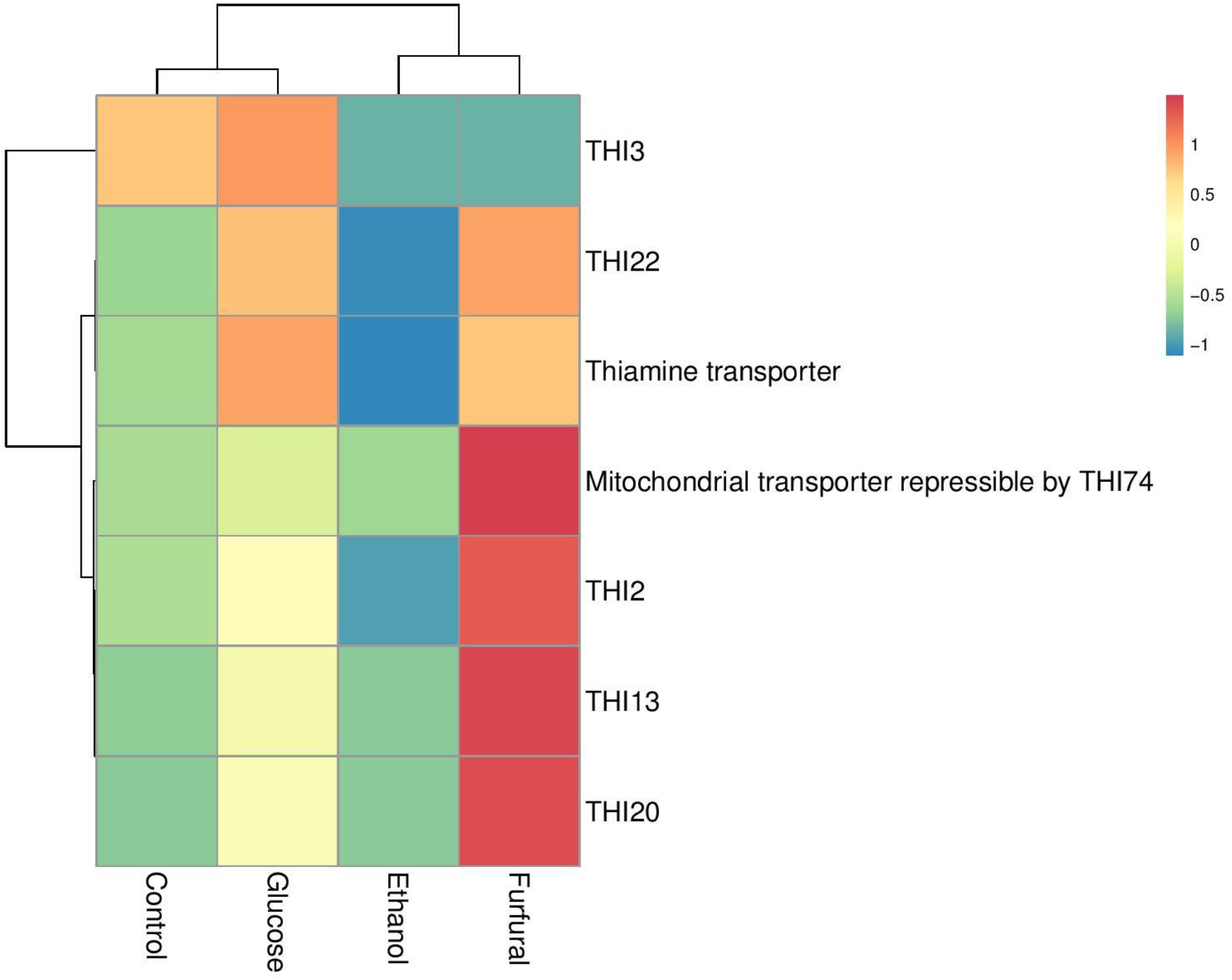
Differential expression of thiamine biosynthesis genes under furfural stress. THI13, THI22, THI2, THI3, THI74, THI20 and a thiamine transporter showed significant and high level of differential expression when treated with furfural

#### 3.4.3. Significant differential expression of various stress-related genes

As ethanol and furfural are potent stress causing agents for the growth and viability of the yeast cells, several stress related genes were among significantly enriched DEGs. Fig. 9 shows that the heat shock proteins HSP26 and HSP12, SSA3, fermentome gene TPS2, hexose transporter HXT7, hexokinase, oxidative stress, osmotic stress genes like SOD2, STR3, GRE2, GLR1, phosphate-sensing TF PHO4, ammonium permeases MEP1,3, mismatch repair protein MLH3, glucose-sensing factor SNF1, stress responsive MSN2, serine/threonine proteins ATG1, GLC7 are some of the important stress induced genes which showed high differential expression levels under stress with respect to control.

**Fig. 9.**
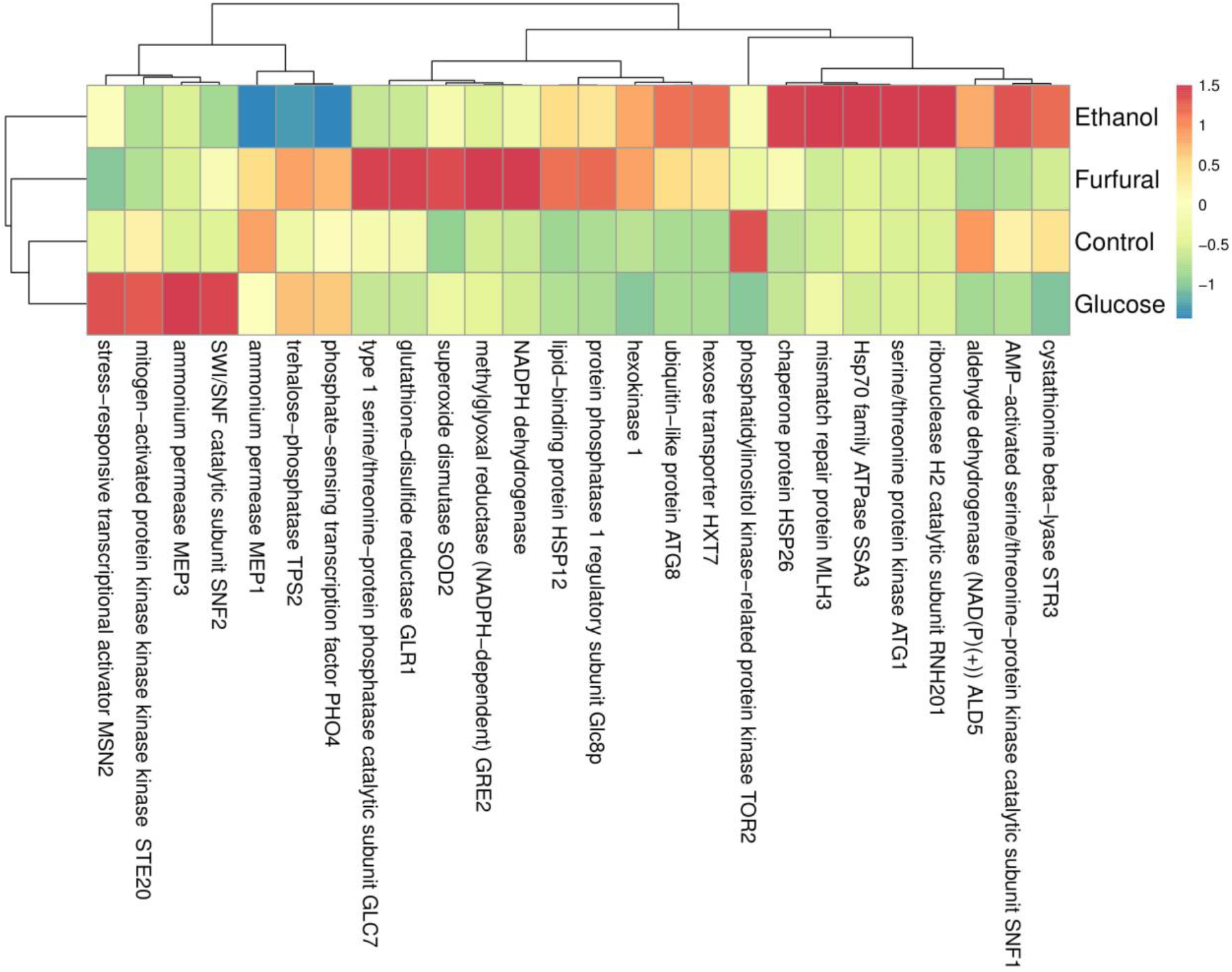
Differential expression of stress associated genes under stress conditions in NCIM3186. Stress associated genes, Hsp26, Str3, Sod2, Hxt7, Ssa3,Tps2, Hxk1, Hsp12 were highly differentially expressed under stress conditions in NCIM3186.

#### 3.4.4. DEGs Annotation, GO ontology and pathway enrichment

Blast2go annotation of the differentially expressed genes resulted in 525 protein-coding genes of which 16 are transcription and translation related proteins, 16 are transporters and 202 are enzymes. Forty eight genes are of non-coding RNA transcripts which include 4 snoRNA, 2 snRNA, 7 tRNA, 35 rRNA coding genes as shown in Fig. 14. A total of 499 GO terms could be enriched within these DEGs by GO annotation tool of blast2go. Fig. 10,11 shows categorization and annotation of GO’s into 3 different components, Biological Process (BP), Molecular Function (MF) and Cellular Component (CC). Interproscan analysis revealed 29 interproscan domains and 19 interproscan families as shown in Fig. 12,13 respectively. KEGG pathway analysis showed that the DEGs were enriched into 78 different pathways among which Glycolysis/Gluconeogenesis, Citrate cycle (TCA cycle), Oxidative Phosphorylation, Inositol phosphate metabolism, Starch and Sucrose Metabolism, Thiamine metabolism, Mitophagy, Meiosis, mTOR signaling pathway, MAPK signaling pathway, Ubiquitin mediated proteolysis, Protein processing in endoplasmic reticulum, Ribosome biogenesis in eukaryotes, mRNA surveillance pathway, RNA transport, ABC transporters are highly enriched pathways which are clearly listed in Table 5.

**Fig. 10.**
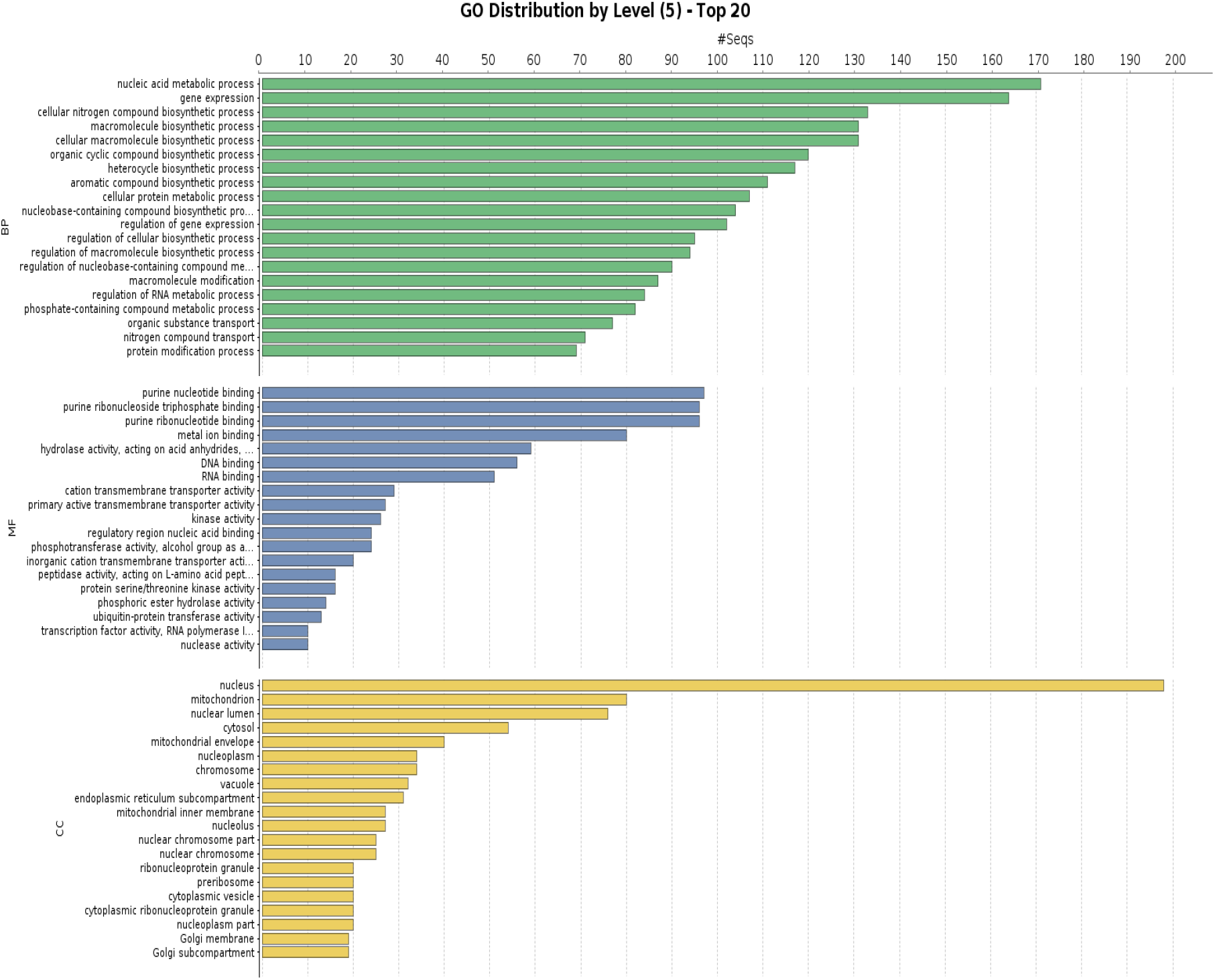
Top 20 Gene Ontology classes enriched by differentially expressed genes in NCIM3186. Nucleic acid metabolic process under biological process, purine nucleotide binding under molecular function and nucleus related processes under cellular component were mostly enriched.

**Fig. 11.**
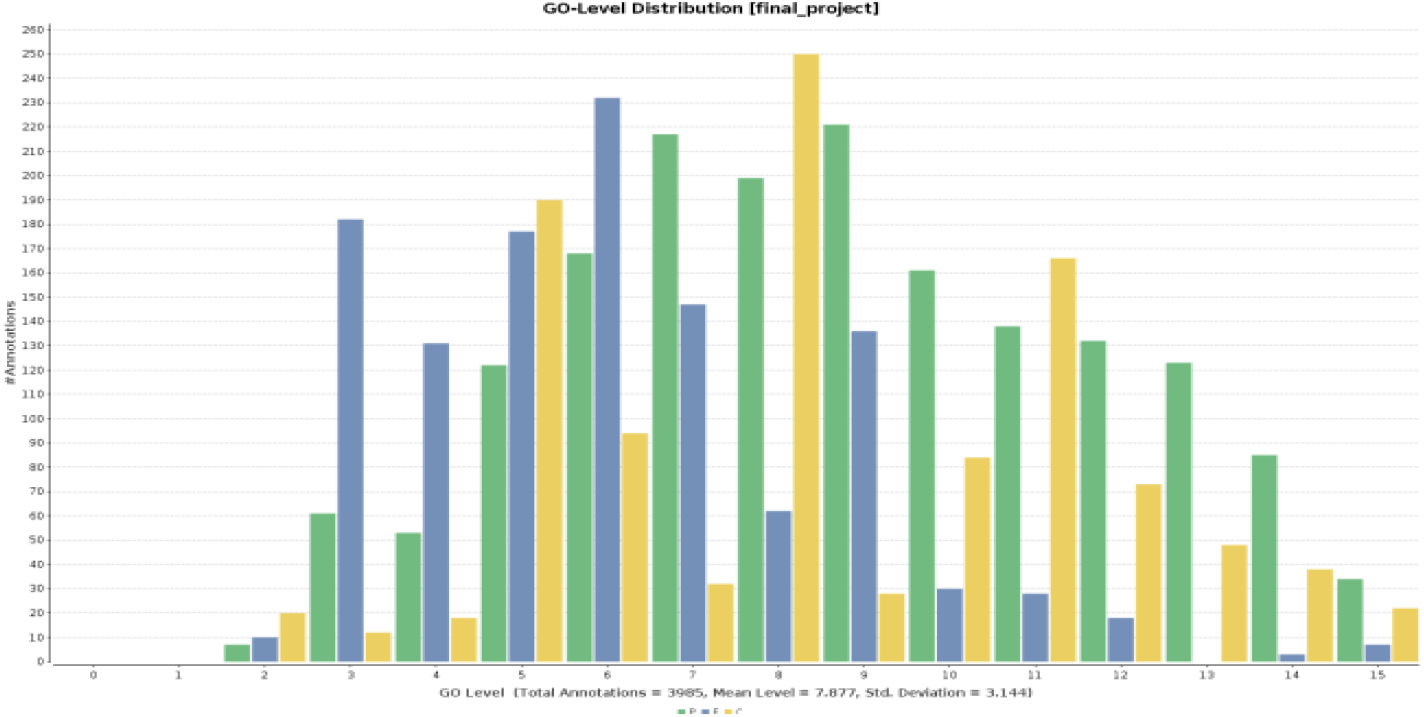
GO-level annotation of differentially expressed genes across the stress treated transcriptomes of NCIM3186. A total of 3985 GOs were annotated from differentially expressed genes of NCIM3186.

**Fig. 12.**
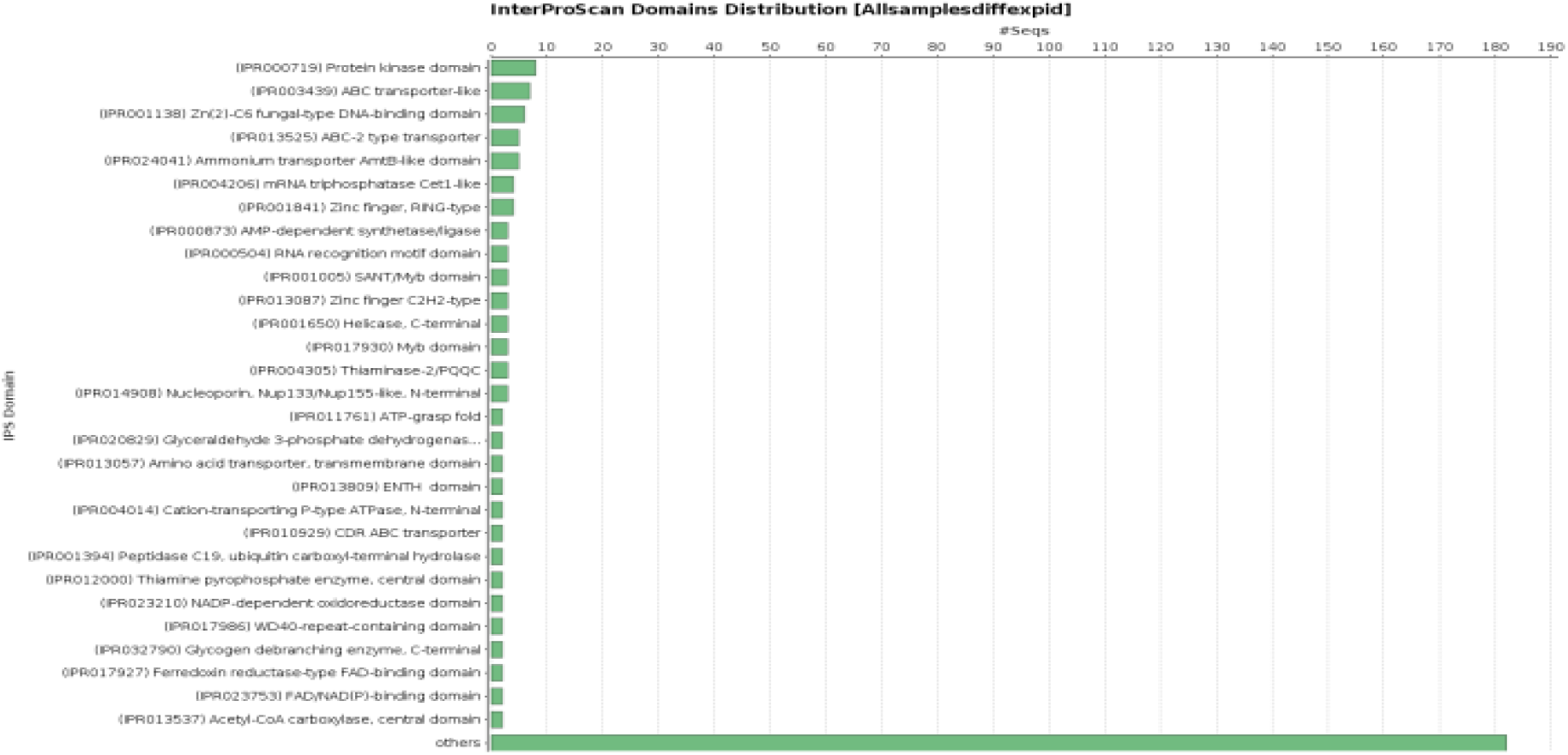
Distribution of InterProScan domains. 29 different interproscan domains were found among the differentially expressed genes in NCIM3186.

**Fig. 13.**
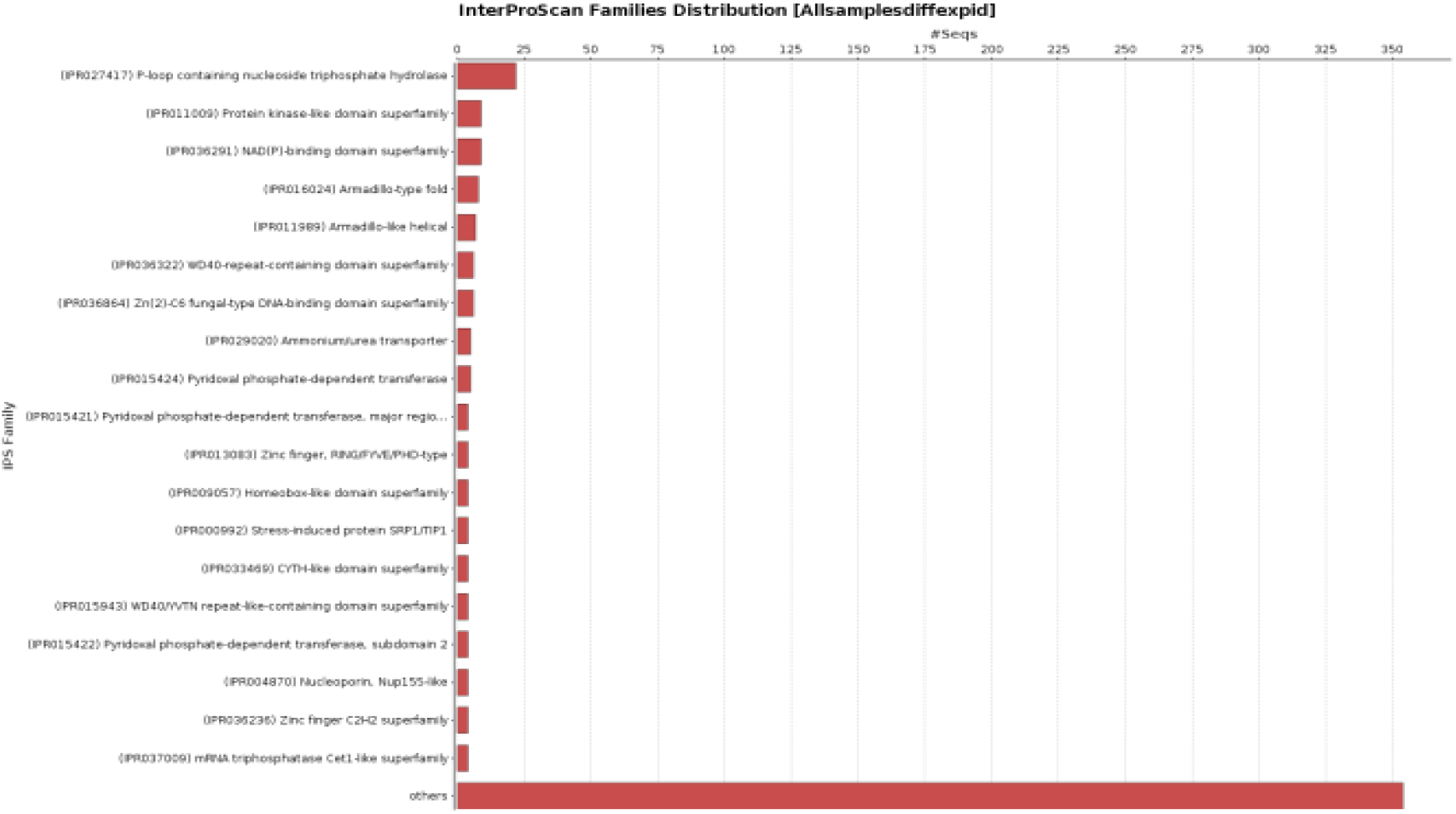
Distribution of InterProScan families. 19 different families were found across the differentially expressed genes in NCIM3186 under stress conditions.

**Fig. 14.**
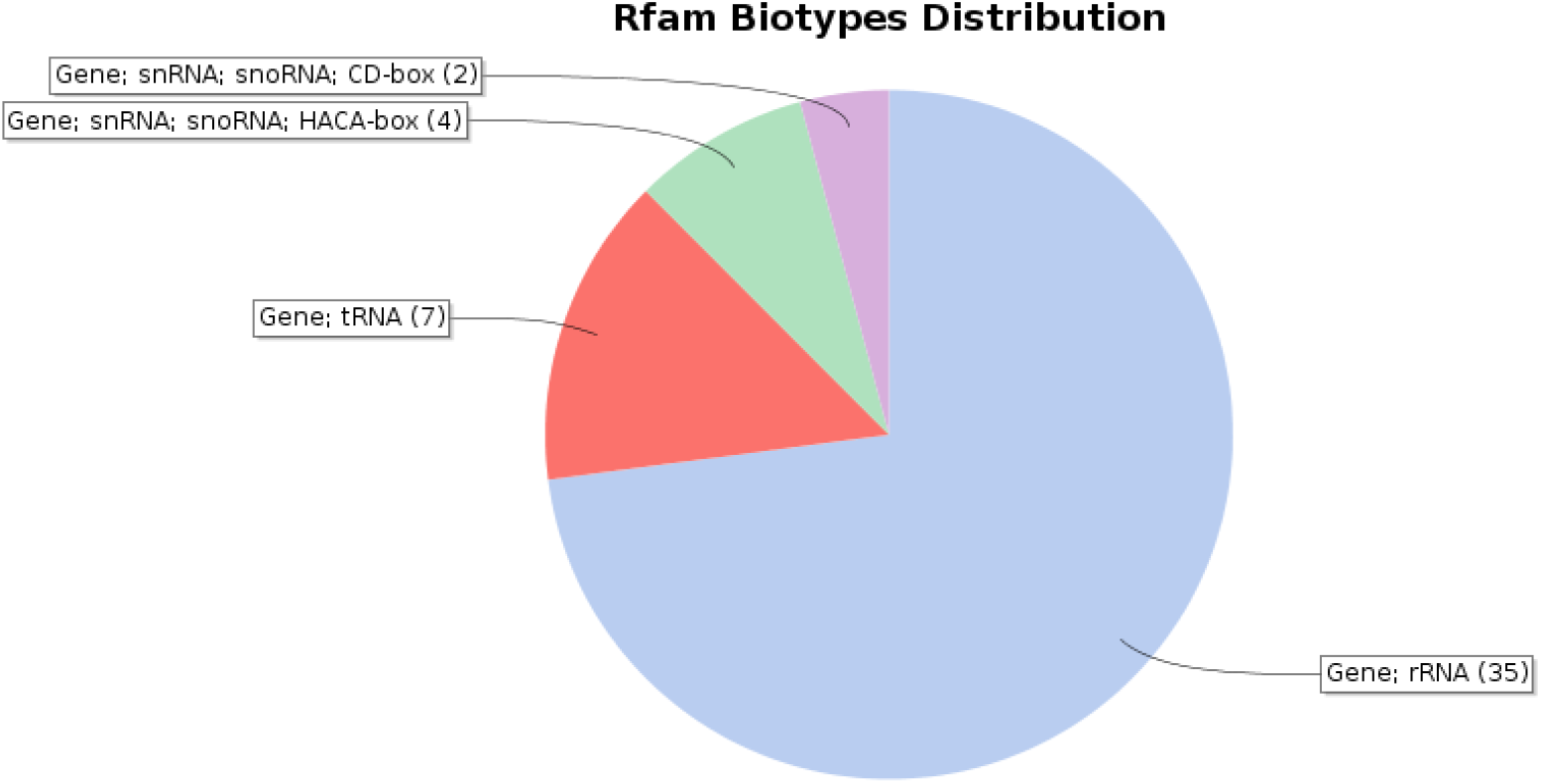
Differentially expressed non-coding RNA genes. Non-coding RNA snRNA, snoRNA, rRNA, and tRNA were differentially expressed in NCIM3186 under stress conditions.

**Table 5.**
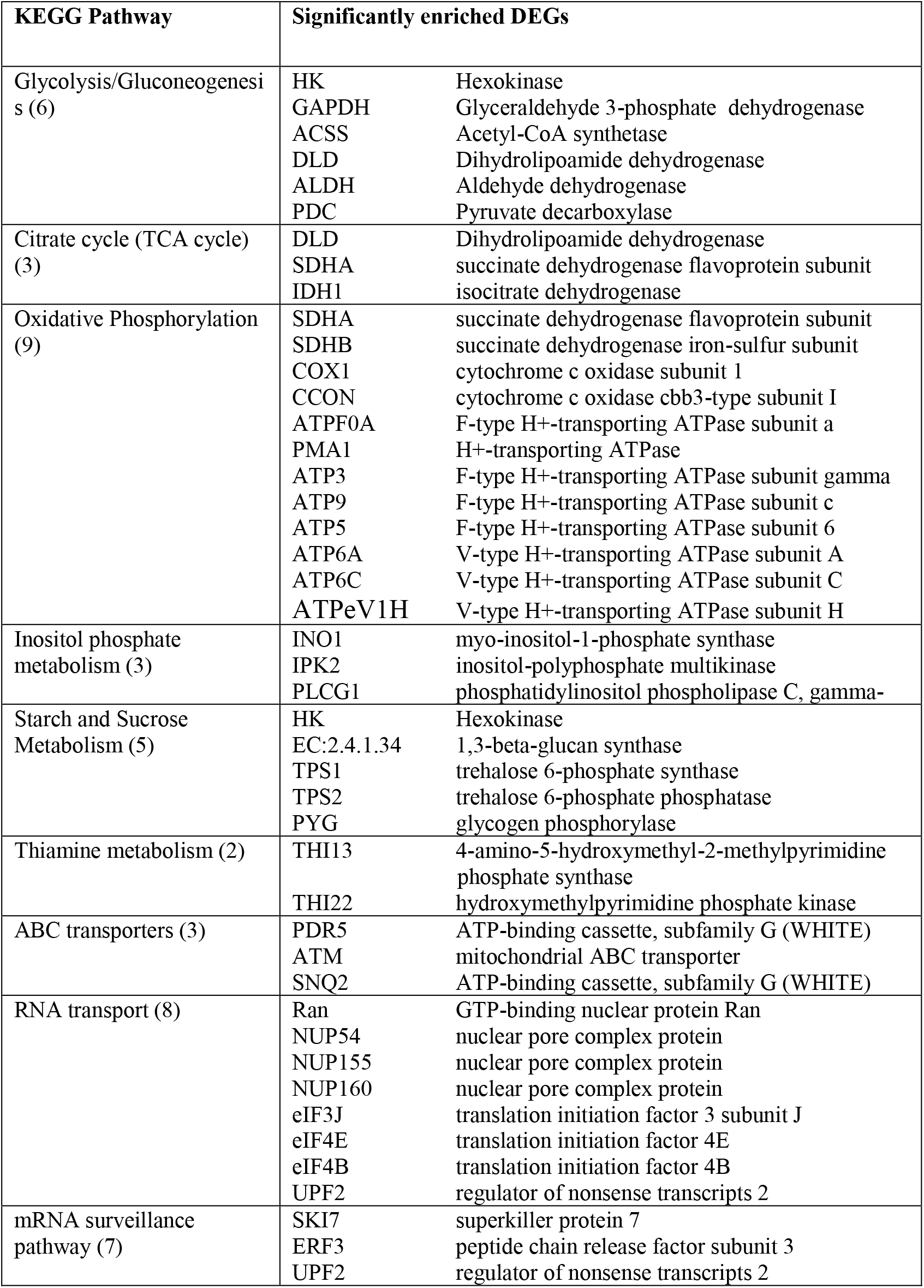
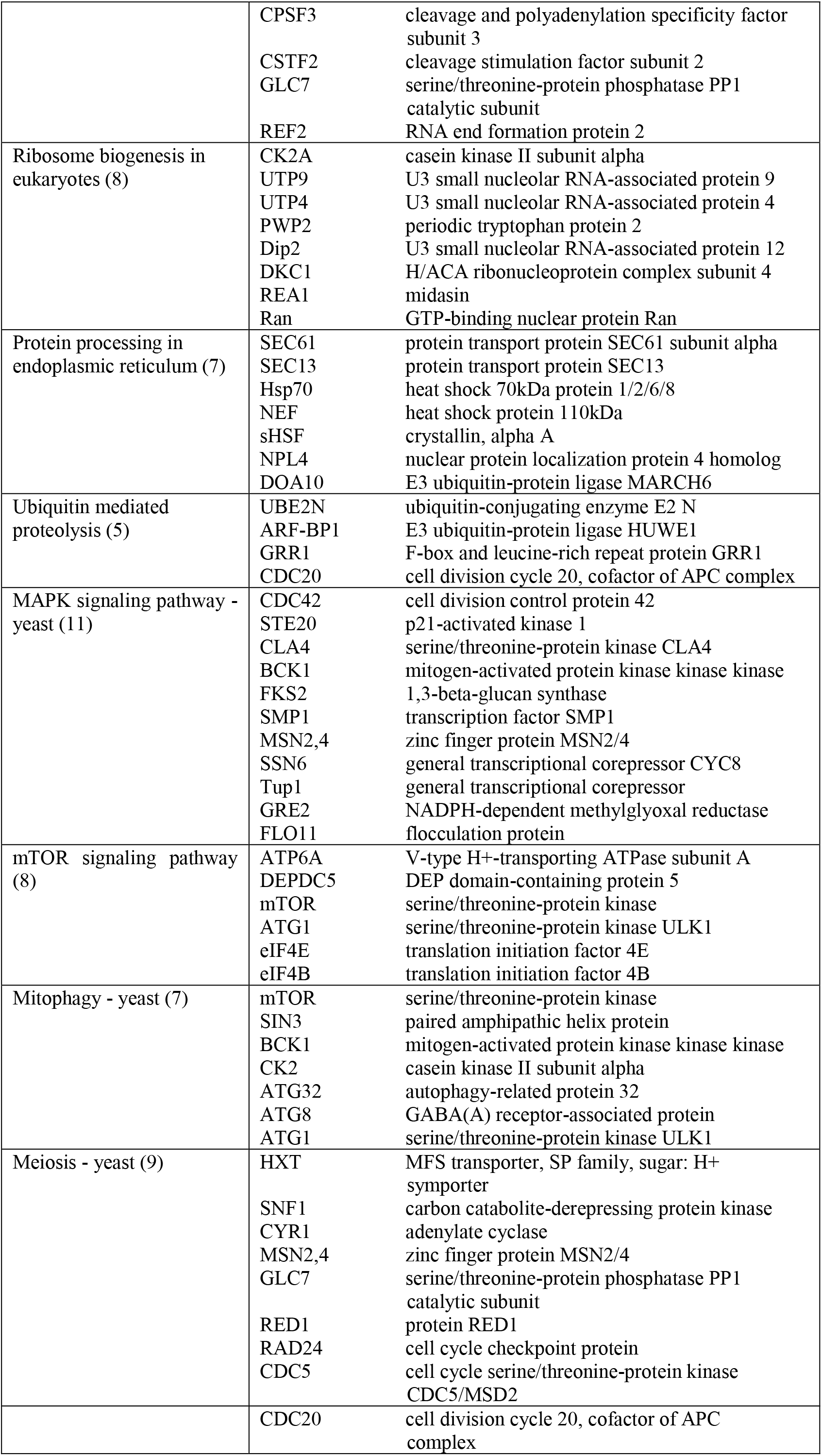
KEGG pathways of significantly enriched differentially expressed genes in NCIM3186

### 3.5. RT-PCR Validation of DEGs

Real time PCR was done to validate the identified DEGs by which we could confirm 6 DEGs. GAPDH was used as a reference gene to normalize the expression of these DEGs. TDH1, HXT6, THI13, TAR1 and 2 other unknown genes (UK-1, UK-2) were the successfully validated DEGs as shown in Fig. 15. TDH1 and HXT6 were highly up-regulated under ethanol stress compared to other conditions which depicts the importance of glycolysis and hexose transporters for maintaining the fermentative behaviour of yeast under a potent solvent stress like ethanol. THI13 and UK-1 are up regulated under furfural and glucose stresses indicating the requirement of free thiamine under inhibitor and sugar stress. HSP12 and UK-2 are highly up-regulated under glucose stress. Most importantly, TAR1 protein was down-regulated under all three stress conditions compared to the control. This underlines that certain genes are even down regulated in yeast in order to maintain the intracellular homeostasis under challenged conditions.

**Fig. 15.**
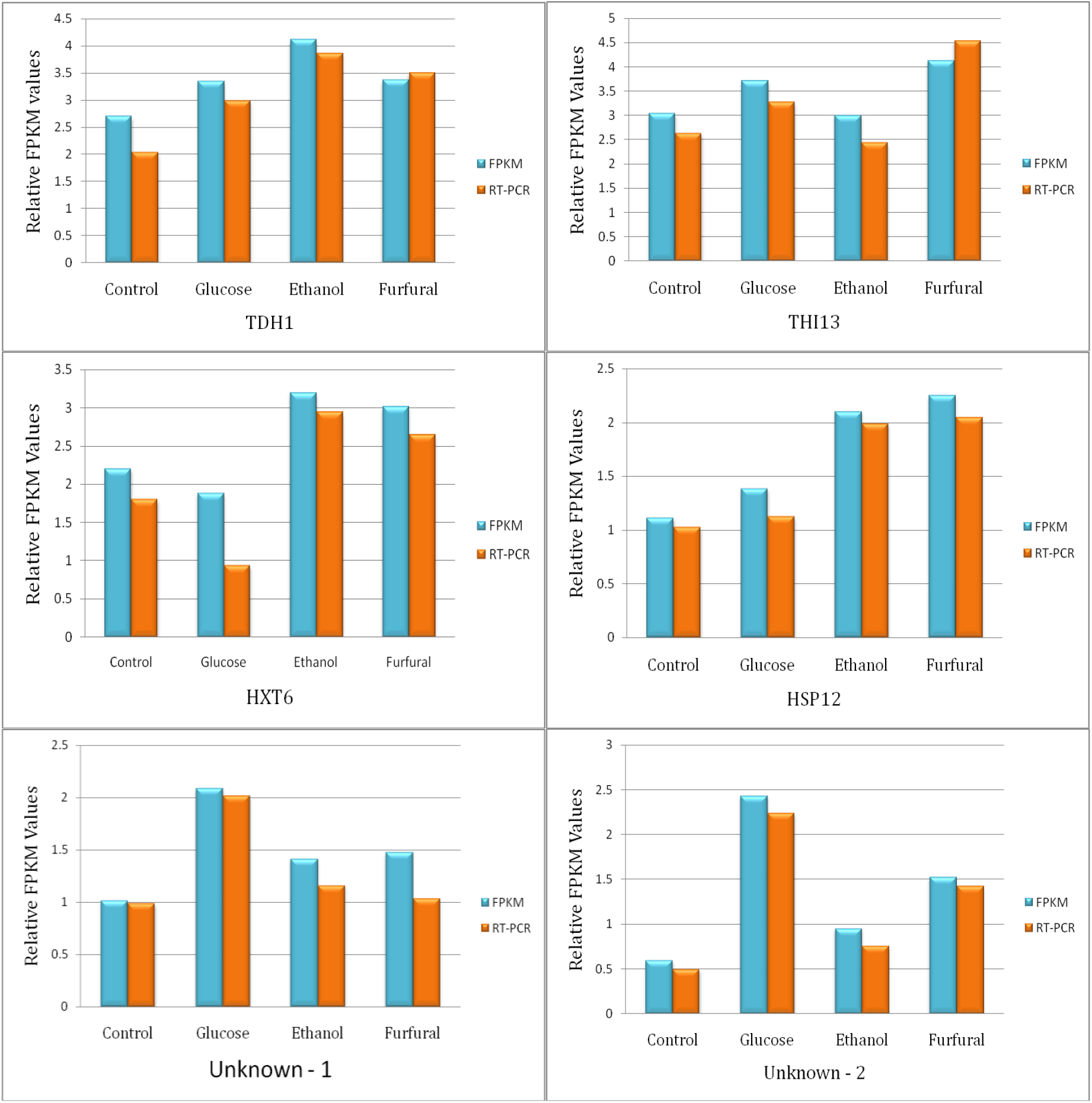
Real-time PCR based confirmation of differentially expressed genes. Differentially expressed TDH1, THI13, HXT6, HSP12, UK-1, UK-2 genes under stress-treated conditions were confirmed based on quantitative real-time PCR analysis.

## 4. Conclusions

RNA-seq based analysis of transcriptomes of NCIM3186 treated with ethanol, glucose and furfural stresses under anaerobic conditions has revealed the expression of a total of 15133 transcripts excluding isoforms. TDH1, TDH2, TDH3, THI13, THI4, CCW12, TAR1, CDC19, snoRNA SNR37, snRNA LSR1, ribosomal proteins P2B, Rpl10, nuclear RNA TPA, non-coding RNA SCR1 were highly expressed genes across the transcriptomes. SCR1, a noncoding RNA was highly expressed gene among all the other genes which was an important observation to be archived. This suggests a regulatory role for non-coding RNAs in yeast cell during expression under stress conditions. A total of 573 genes were differentially expressed at P_value_ = 1e-3 and fold change of 2. TDH1, THI13, HXT6, PDR5, HSP26, HSP12, STR3, INO1, TAR1, SSA3, MNT3, PEX6, RGI1, IRC8, VMA13, FAA4, YRO2, OLI1, intron-encoded reverse transcriptase al2 and 2 other unknown proteins showed significant differential expression. Down regulation of TAR1 gene under stress conditions in comparison to control transcriptome was found to be an interesting observation in NCIM3186 expression pattern under stress.

Up regulation of thiamine biosynthesis pathway genes, THI13, THI2, THI3, THI20, THI22, THI74 repressible mitochondrial transporter and a thiamine transporter under furfural stress has evidently shown that there exists a relationship between thiamine and yeast cellular stress responses. Stress associated genes, HSP26, HSP12, SSA3, fermentome gene TPS2, HXT7, hexokinase, oxidative stress, osmotic stress genes like SOD2, STR3, GRE2, GLR1, phosphate-sensing TF PHO4, ammonium permeases MEP1,3, mismatch repair protein MLH3, glucose-sensing factor SNF1, stress responsive MSN2, serine/threonine proteins ATG1,GLC7 were differentially expressed under stress in NCIM3186. This work claims to be the first RNA sequencing based study to analyze the differential response of yeast when treated with ethanol, glucose and furfural under anaerobic conditions. Thus, our study proves to be an ultimate and promising way to uncover yeast gene expression patterns under stress conditions with the aid of high-throughput sequencing technologies to serve the purpose of RNA sequencing unlike earlier traditional microarray based studies.

## Abbreviations

DEG-: Differentially expressed gene
YEPD-: Yeast extract peptone dextrose
TPM-: Transcripts per million
TMM-: Trimmed mean of M-values
FPKM-: Fragments per Kilobase of Exon per Million Fragments Mapped transcripts per million
snRNA-: small nuclear RNA
snoRNA-: small nucleolar RNA
rRNA-: ribosomal RNA
tRNA-: transfer RNA
KEGG-: Kyoto Encyclopedia of Genes and Genomes
KAAS-: KEGG Automatic Annotation Server
BP-: Biological Process
MF-: Molecular Function
CC-: Cellular Component

## Acknowledgements

Financial Assistance from University Grants Commission (OU-UGC-UPE) under the University of Potential for Excellence programme is acknowledged. B. Sravanthi Goud thanks the University Grants Commission for the BSR-RFSMS Senior Research Fellowship.

